# UDP-glucose dehydrogenase Ugd in *E. coli* is activated by Gmd and RffD, is inhibited by CheY, and regulates swarming

**DOI:** 10.1101/2020.01.08.899336

**Authors:** Irina A. Rodionova, Zhongge Zhang, Mohammad Aboulwafa, Milton H. Saier

**Author notes:** To whom correspondence should be addressed: Milton H. Saier Jr: Department of Molecular Biology, Division of Biological Sciences, University of California at San Diego, La Jolla, CA 92093-0116, USA.

## Abstract

The two most common mechanisms of polymyxin resistance in bacteria involve glycosylation of the outer membrane lipopolysaccharide (LPS) and production of the exocapsular polysaccharide, colanic acid (CA). UDP-glucose dehydrogenase, Ugd, is required for both CA biosynthesis and LPS modification. We here show that Ugd is activated by the GDP-mannose-4,6-dehydratase (Gmd, YefA, YefN), UDP-N-acetyl-D-mannosamine dehydrogenase (RffD, WecC), and ribonuclease HII (RnhB). The former two enzymes are involved in Lipid A and colanic acid biosyntheses, respectively, while RnhB cleaves RNA in RNA:DNA hybrids. Moreover, CheY inhibits the phosphorylated, activated form of Ugd (Ugd-P). Finally, Ugd is involved in the regulation of swarming, since a *ugd* mutant has an increased swarming rate, while Ugd overproduction inhibits swarming. Two-hybrid bacterial assays reveal direct interaction of Ugd with RssB (an anti-RpoS factor) and CheY in vivo.

## Introduction

UDP-glucose dehydrogenase, Ugd, catalyzes the oxidation of UDP-glucose, yielding UDP-glucuronate, coincident with the reduction of NAD, and this reaction is essential for colonic acid (CA) biosynthesis. Ugd is conserved in many bacteria and is transcriptionally regulated as part of the RcsB, PhoP, and PmrA regulons in *E. coli* (1). Thus, the participation of different combinations of regulatory systems indicate that *ugd* expression is responsive to a variety of environmental and internal signals, including changes in temperature, osmolarity, polymyxin B concentration and iron concentration (1,2). Moreover, the presence of bile salts induces resistance to polymyxin B in enterohemorrhagic *E. coli* (3). Nevertheless, polymyxin is an effective drug for *mcr-1* antibiotic resistant *E. coli* (4), and polymyxin combinations combat *E. coli* harboring *mcr-1*, as well as blaNDM-5, and therefore provides promise for postantibiotic development (5).

Ugd, as well as the uridylyltransferase, GlnD, and the GTPase, MnmE, activate folylpoly-gamma-glutamate synthase, FolC in *E. coli*, by direct protein-protein interactions (6). Thus, Ugd appears to be part of a complex regulatory system, influencing and influenced by many types of physiological states and conditions.

The operon for colanic acid biosynthesis includes the *wzaABC-wcsABCDEF-gmd-fci-gmm-wcaI-manCB-wcaJ-wzx-wcaKLM* genes (7). The *ugd* gene is located separately in the *E. coli* genome. Ugd, mildly activated by ATP at inhibitory NAD concentrations (8), is also involved in the production of UDP-4-amino-4-deoxy-L-arabinose, a compound that modifies Lipid A, rendering *E. coli* resistant to cationic antimicrobial peptides such as polymyxin B (2,9). Ugd activity is regulated by tyrosyl phosphorylation, catalyzed by the tyrosyl kinase, Wzc (10), and the phosphorylation of Ugd regulates the production of colanic acid (2). Phosphorylation of Ugd by Wzc has been found to participate in the regulation of the amount of the colanic acid (CA) exopolysaccharide, whereas Etk-mediated Ugd phosphorylation appeared to promote resistance of *E. coli* to polymyxin (2).

Allosteric regulation of Ugd activity has been reported before (8), but protein-protein interactions represent additional potential mechanisms for Ugd regulation by proteins related to exopolysaccharide biosynthesis and envelope stress. We have found that Ugd interacts with the chemotaxis regulator, CheY, which transmits a signal to the flagellar motor component, FlgI, a flagellar basal body protein. In fact, Ugd interacts with RnhB (ribonuclease HII that degrades the RNA in DNA-RNA hybrids), and RnhB is conserved in a gene cluster with the genes encoding LpxA and LpxB as well as Lipid-A-disaccharide synthase (EC 2.4.1.182), all of which participate in Lipid A biosynthesis. Ugd also interacts with Gmd (GDP-D-mannose dehydratase), which is stabilized by a protein-protein interaction with WcaG (11) and is involved in CA biosynthesis. Finally, Ugd interacts with WcaA, a glycosyl transferase, and with RffD (WecC) (UDP-N-acetyl-D-mannosaminuronic acid dehydrogenase), involved in Enterobacterial Common Antigen biosynthesis (12,13). The interaction scores of Ugd with these and other proteins are presented in Table 1 (see the BioGrid database https://thebiogrid.org/4261356). The method that was used for the detection of protein interactions with Ugd has been described (14).

**Table 1.**
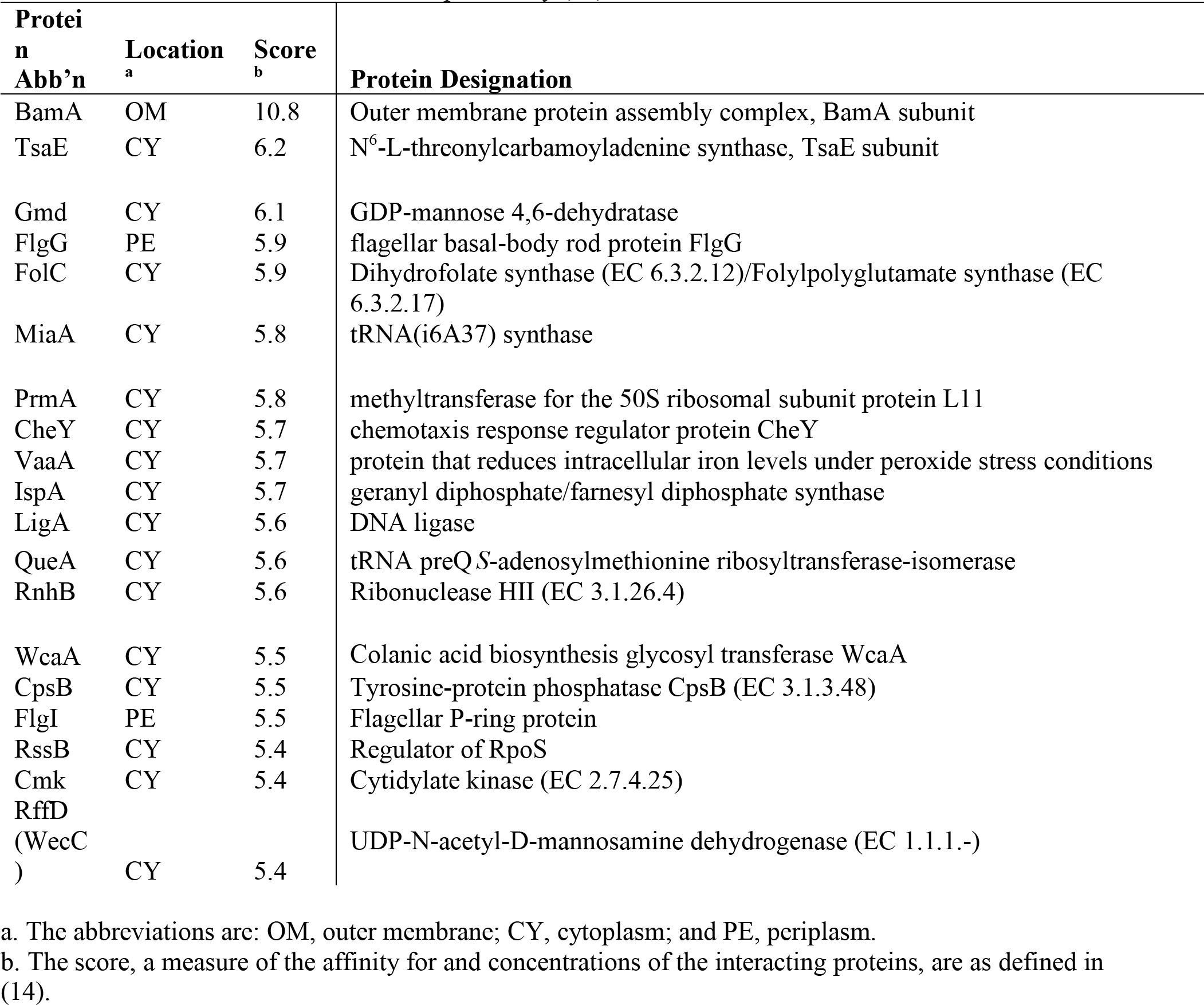
Interactions of Ugd with other cellular proteins in *E. coli* and determination of the interaction scores which were measured as described previously (14).

The Wzc protein has a cytoplasmic tyrosyl phosphorylation domain that activates Ugd by phosphorylation, producing Ugd-P, dependent on autophosphorylation of Wzc on tyrosine 569 (10). Wzc is also a polysaccharide co-polymerase protein with two transmembrane segments and a large periplasmic domain that interacts with the outer membrane channel, Wza. Considering the possible regulation of Ugd by CheY, we tested Ugd-P activity in the presence and absence of the CheY protein and observed strong inhibition. Moreover, we show an increased swarming rate for a Δ*ugd* mutant, relative to parental control strains, and overexpression of *ugd* depressed swarming.

As noted above, CheY is a regulator of flagellar rotation, and it binds to the flagellar rotor in response to concentration gradients of chemoattractants and repellants. The *E. coli* CheY modulates the rotational bias of the flagellar motor, which controls the run-and-tumble motion of the bacterium, allowing it to migrate towards regions of high chemoattractant concentrations and away from high concentrations of repellants. Thus, the switching mechanism of the rotational direction determines the runs and tumbles of the cell in a way dependent on the CheY-P level (15).

CheY protein-protein interactions have been detected with the flagellar membrane protein complex including FliGHMN, FlgGI and FlhA (14). A stochastic conformational spread model has been invoked to explain the dynamic and cooperative behavior of the bacterial flagellar motor (16). Chemotactic adaptation of *Yersinia pestis* within neutrophils is mediated by CheY, which in this organism, not only controls the direction of flagellar rotation, but also upregulates the modification of lipid A (17,18). Also, analysis of a *Δugd* mutant of *E. tarda* revealed a truncated core oligosaccharide region of LPS. The mutant exhibited enhanced autoaggregation and biofilm formation as well as reduced hemolytic activity (19).

CA confers the ability to survive at high temperatures and in the presence of bile salts, mainly by enhancing biofilm formation. CA in *E. coli* has been reported to bolster biofilm formation by maintaining the bacterial interaction and biofilm three-dimensional structure, thus preventing swarming. Genes essential for motility and swarming have been identified by screening the *E. coli* non-essential gene mutant collection (20). These genes include cell envelope and outer membrane biogenesis genes as well as those concerned with cell motility, signal transduction and lipid metabolism, but *ugd* and *wzc* mutants were not identified as influencing swarming (20).

Interaction of CheY with Ugd in *E. coli* has been suggested (14) and confirmed in this paper using a bacterial two-hybrid assay. We further demonstrate that Ugd is allosterically activated by the WecC and RnhB proteins in the presence, but not in the absence, of ATP. However, for Gmd-dependent Ugd activation, the presence of ATP proved not to be required. These observations indicate how CA production may be regulated in response to defects in the production of other exopolysaccharides, detected in *E. coli* mutants of *waaF*, *waaG* and *waaP* (7). A compensatory mechanism of Ugd regulation may explain why CA was overproduced when production of another exopolysaccharide was prevented (7).

## Experimental Procedures

### Protein purification

The overproducing strains for the proteins Ugd, WecC, CheY, FolC and RnhB from the Keio collection were grown in LB medium (1 L), induced by addition of 0.5 mM IPTG when the culture reached an OD_600_ = 0.8, and cells were harvested after 4 h of shaking at 250 rpm. Then, the cells were resuspended in 20 mM HEPES buffer, pH 7, containing 100 mM NaCl, 2 mM β-mercaptoethanol and 0.3 % Brij 30 with 2 mM phenylmethylsulfonyl fluoride present. Cells were lysed by incubation with lysozyme (1 mg/ml) for 40 min at 4°C, followed by a freeze-thaw cycle and sonication. The pellets were collected after lysis, sonication, and centrifugation at 14,000 rpm for 25 min, and the insoluble fraction was resuspended in At-buffer containing 7M urea and 0.3% Brij 30. Inclusion bodies were dissolved, and after sonication and centrifugation, they were analyzed. The RnhB protein was refolded from the insoluble fraction as described in (1). Protein size, expression level, distribution between soluble and insoluble forms, and extent of purification were monitored by SDS-PAGE. Protein concentrations were measured using the Bradford assay kit (Bio-Rad).

### Regulation of Ugd-P activity by direct protein-protein interaction with CheY

The standard enzymatic assay applied to the conversion of UDP-glucose to UDP-glucuronate, catalyzed by 200 nM Ugd, results in the reduction of NAD accompanied by an increase in absorbance at 340 nm. The assay mixture for kinetic measurements included: 100 mM Tris-glycine, pH 8.3, 0.1 M NaCl, 5% glycerol, 2 mM DTT, 1 mM ATP, 2 mM NAD and 0 – 6 mM UDP-glucose. Ugd was preincubated with Wzc for 30 min at 30 °C. The activity was measured in the presence and absence of 500 nM CheY. Phosphorylation of Ugd by the tyrosine kinase domain of the Wzc transport auxillary protein has been investigated by Grangeasse who kindly provided the overexpression construct for the Wzc kinase domain for the phosphorylation of Ugd (Ugd-P) (10).

### Regulation of Ugd activity by direct protein-protein interactions with RnhB and WecC

The Ugd activity was measured at 0-5 mM NAD concentrations, and no inhibition was found under these assay conditions. Enzyme assays, in which NADH was detected in the assay mixtures, included Tris-glycine buffer, pH 8.3, glycerol, 2.5%, 0.1 M NaCl, 2 mM NAD, 5 mM ATP, 2 mM DTT and variable concentrations of UDP-glucose. The Ugd kinetic parameters were calculated using Prism 7 (GraphPad Prism).

### Regulation of Ugd activity by direct protein-protein interaction with Gmd

The assay mixture included 0.1 M Tris buffer, pH 8, 2.5% glycerol, 0.1 M NaCl, 1 mM NAD, 0-0.65 mM ATP, and 0.7 mM UDP-glucose.

### Constructions of bacterial two-hybrid plasmids

The Euromedex bacterial two-hybrid system (Euromedex, Cat # EUK001), consisting of two compatible plasmids pUT18 and pKNT25, is designed to detect and characterize protein-protein interactions *in vivo*. These two plasmids encode the T18 domain and the T25 domain of cAMP synthase of *Bordetella pertussis*, respectively. To determine any possible interactions between two target proteins, one gene is fused to the N-terminus of T18 in pUT18 while the other is fused to the N-terminus of T25 of pKNT25. These two plasmids can be co-transformed to an *E. coli* strain lacking its own cAMP synthase gene *cyaA*. In the case that the two target proteins closely interact, thereby bringing together the T18 and T25 domains, the *E. coli* Δ*cyaA* cells enable synthesis of cAMP, thereby exhibiting unique phenotypes that can be quantitated.

The *cheY* structural gene with no stop codon was amplified by PCR from BW25113 genomic DNA using carefully designed oligonucleotides (Table S2), digested by *Hind*III and *BamH*I, and then ligated into the same sites of pUT18 individually. In each of these resultant plasmids, the target structural gene (with no stop codon) is located immediately upstream of the N-terminus of T18 in pUT18, creating a single hybrid gene that encodes a two-domain protein. This plasmid is denoted pUT18-*cheY* (Table S1).

Similarly, the *ugd* structural gene with no stop codon was amplified, digested with *Hind*III and *BamH*I, and ligated into the same sites of pKNT25. The resultant plasmid carries the *ugd* gene, in frame fused to the N-terminus of the T25 domain. It was denoted as pKNT25-*ugd.*

To test the possible interaction between *ugd* and *cheY* and between *ugd* and *rssB*, plasmid pairs of pKENT25-*ugd*/pUT18-*cheY* and pKENT25-*ugd*/pUT18-*rssB* were individually co-transformed by electroporation into the *E. coli* BTH101 strain deleted for *cyaA*. The Km^r^ and Ap^r^ transformants were inoculated onto MacConkey agar plates supplemented with maltose, Km, Ap and IPTG. The plates were incubated at 30 °C for up to 48 h. The appearance of red colonies showed positive interactions between the target proteins. The intensity of red color proportionally correlates with the amount of the protein-protein interactions.

## Results

Ugd of *E. coli* is an enzyme responding to various stresses, such as the presence of polymyxin B. Ugd is encoded separately from the CA biosynthetic operon, not only in *E. coli*, but also in the genomes of many Proteobacteria, and in those organisms that have been examined, it is also overproduced in response to multiple stress conditions (1). Purified WecC, CheY, RssB, FolC and RnhB proteins, involved in Ugd interactions (Table 2) (14), were tested for regulation of Ugd activity in the presence and absence of ATP. Phosphofructokinase (PfkB), previously characterised (21), was used as a negative control.

**Table 2.**
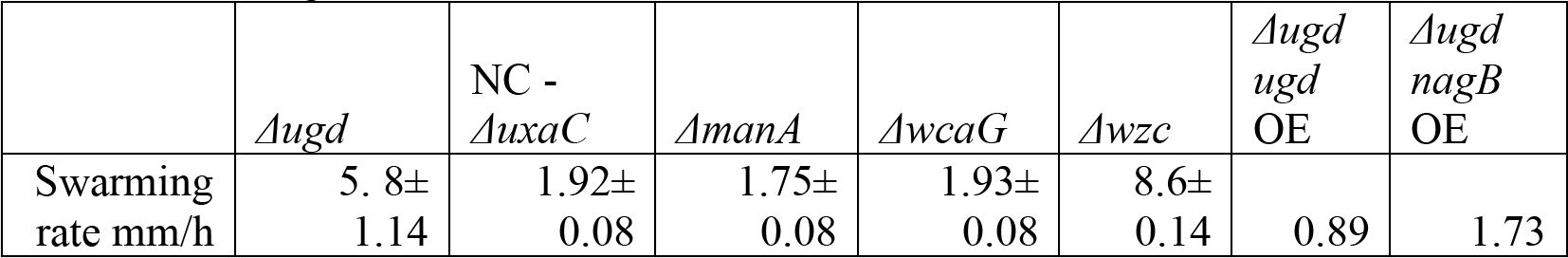
Swarming rates for mutants in *E. coli* (Keio collection).

### Inhibition of Ugd by CheY

The allosteric inhibitory effect of CheY on Ugd-P is presented in Fig. 2. The steady-state kinetics for Ugd-P activity were measured in the presence and absence of CheY. CheY was shown to substantially inhibit the activated Ugd-P using the soluble Wzc domain for Ugd phosphorylation. CheY was added to the assay mixture at a final concentration 2.5-fold higher that of Ugd. Ugd-P had only 5-10% of its uninhibited activity when CheY was present in this amount (Fig.2).

**Figure 1.**
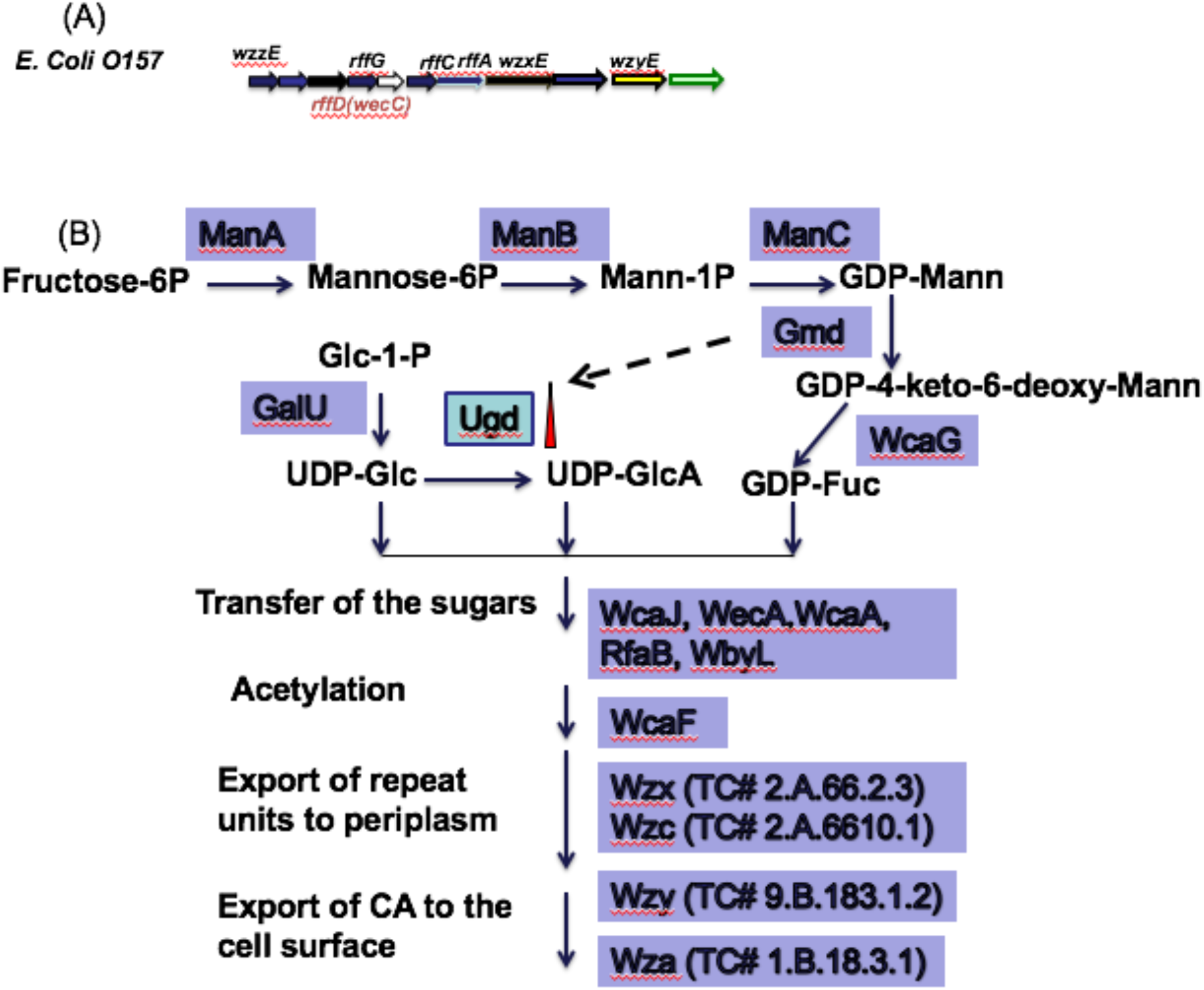
(A) Lipid A operon structure of *E. coli*, including *rffD* (*wecC*). (B) The pathway for CA biosynthesis in *E. coli*, showing the compounds involved (in bold), with the enzymes indicated by the boxed protein abbreviations adjacent to the arrows between the substrates and products. Abbreviations: the mannose-6-phosphate isomerase (ManA), phosphomannomutase (ManB), GDP-D-mannose synthetase (ManC), GDP-mannose dehydratase (Gmd), GDP-L-fucose synthase (WcaG). Other abbreviations can be found in the text or are functionally defined in the figure.

**Figure 2.**
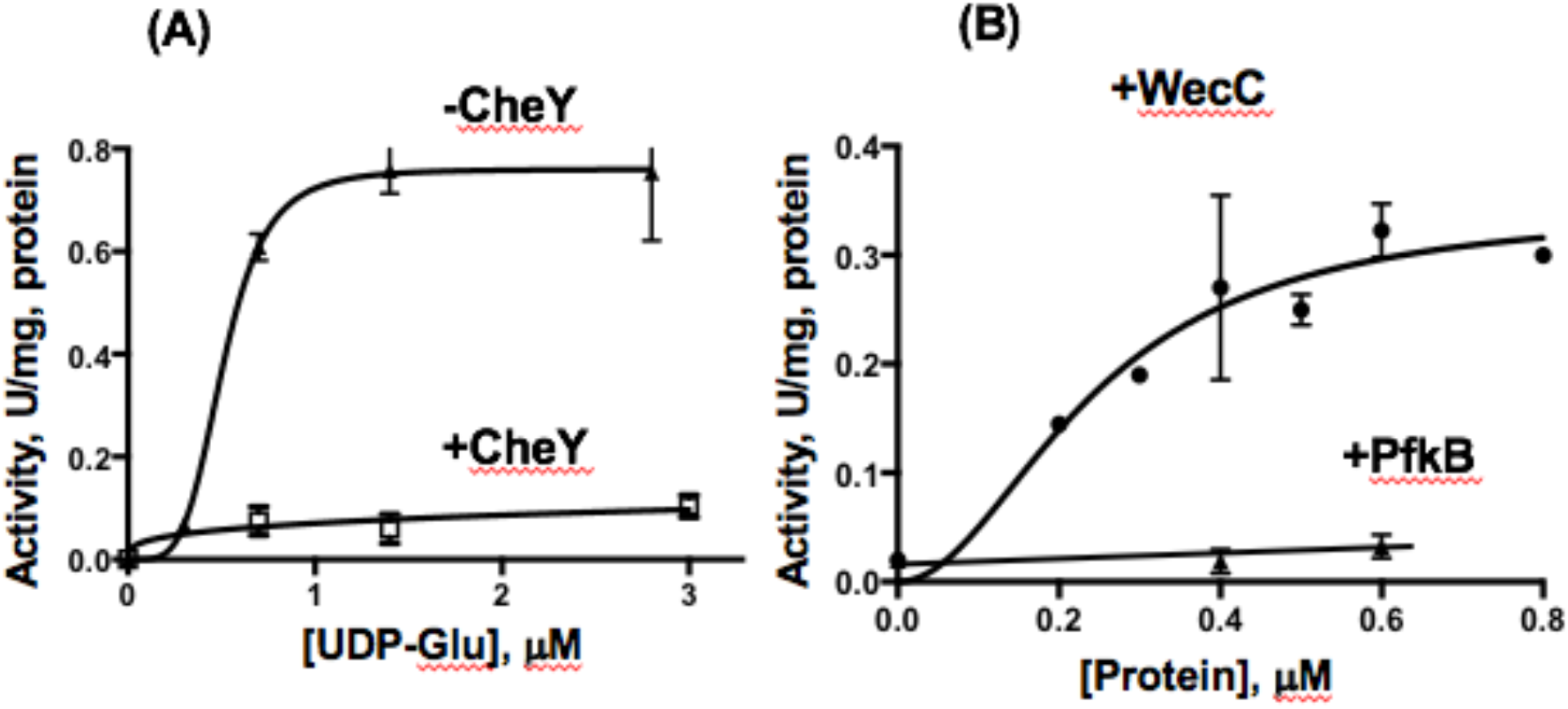
Regulation of Ugd by protein-protein interactions. (A) Allosteric inhibition of Ugd-P by CheY, and (B) activation of Ugd by WecC. (A) The steady-state kinetics of Ugd-P were determined as a function of the UDP-glucose concentration (0 to 3 mM) in the absence of CheY (closed triangles and diamonds) or the presence of CheY (open squares). (B), the activity of Ugd was measured as a function of the WecC concentration, 0-0.8 μM, with 0.5 mM UDP-glucose (circles). PfkB (triangles) was used as a control.

### Ugd activation by RnhB and WecC

We found that Ugd activity is substantially upregulated by RnhB (ribonuclease HII). The control protein, FolC, gave a negative result as neither inhibition nor activation was observed (Fig. 3). In both cases, the activity of Ugd was measured as described under “Experimental Procedures”. Addition of 5 mM ATP to the UDP-glucuronate-producing assay mixture increased the V_max_ 2-fold at a 0.8 mM UDP-glucose concentration.

**Figure 3.**
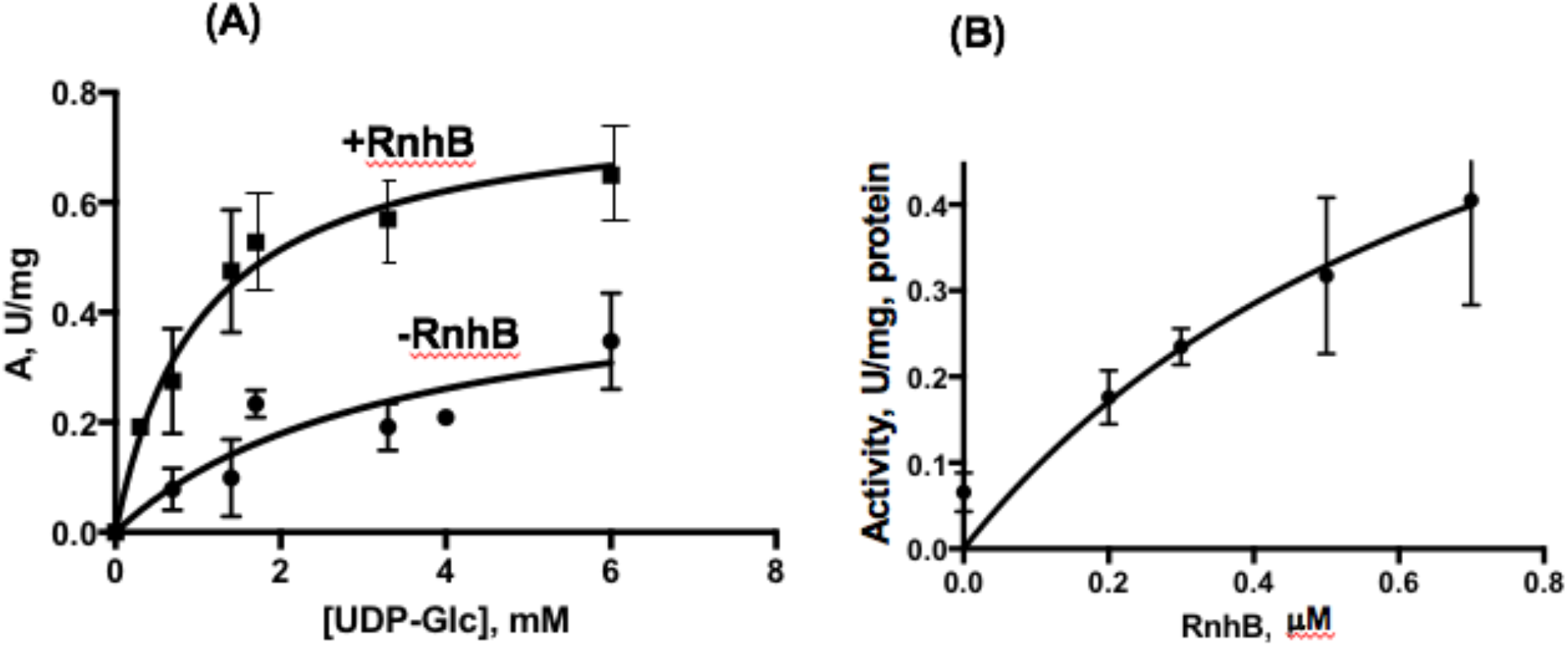
Allosteric activation of Ugd by RnhB. (A), The steady-state kinetics of Ugd-P in the presence (squares) and absence (circles) of RnhB were determined as a function of the UDP-glucose concentration (0 to 6 mM) with 1 mM ATP present in all reaction mixtures. (B). Activation of Ugd as a function of the RnhB concentration when the concentration of UDP-glucose in the assay mixture was 0.7 mM. Units are expressed in μmoles/min/mg, protein.

Titration of the Ugd activation effect of WecC at 0-1 μM concentrations is shown in Fig. 3. In this experiment, the UDP-glucose concentration was 0.6 mM. The calculated K_d_ for the WecC-Ugd interaction was calculated to be 0.24±0.01 μM of WecC. Phosphofructokinase PfkB (21) was used as a negative control, and no activation of Ugd was observed when this protein was added at a concentration of 0.6 μM (Fig.3).

### Ugd activation by Gmd

We found that Ugd activity is substantially upregulated by Gmd. Titration with Gmd is shown in Fig 4A. The absence of ATP corresponds to the zero value in Fig. 4B. The dependency of Gmd-dependent Ugd activation on the ATP concentration is shown in Fig. 4B at 1 mM NAD, 0-0.65 mM ATP, and 0.7 mM UDP-glucose. Activation in the absence of ATP was observed; however, the maximum for the Gmd-dependent activation of Ugd was observed at a 0.1 mM ATP concentration, corresponding to a physiologically relevant concentration in *E. coli*.

**Figure 4.**
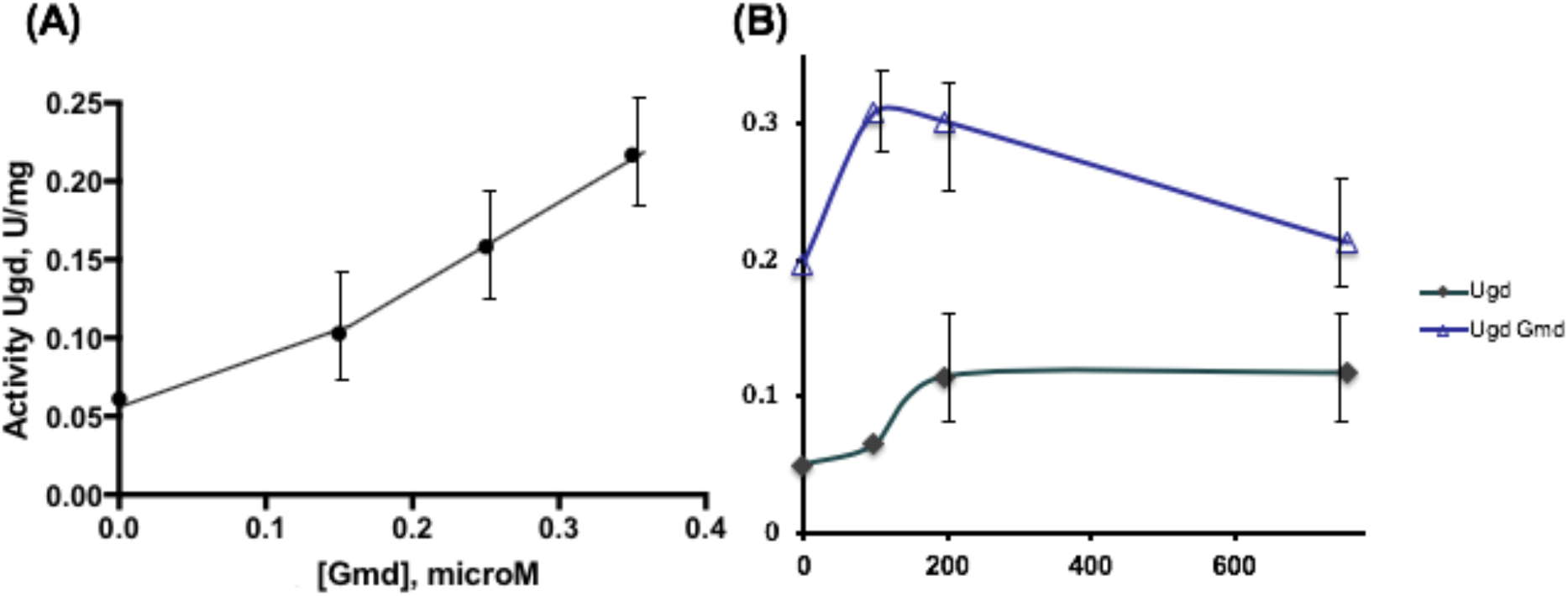
Allosteric activation of Ugd by Gmd. (A), The activity of Ugd was measured as a function of the Gmd concentration, 0-0.4 μM, with 0.5 mM UDP-glucose present (circles). (B), The steady-state kinetics of Ugd were determined as a function of the ATP concentration (0 to 800 μM) in the absence (diamonds) or presence of 0.5 μM Gmd (triangles) at pH 8, 1 mM NAD and 0.7 mM UDP-glucose. Units = μmoles/min/mg, protein.

### Swarming

The assay for swarming motility by various *E. coli* K-12 strains was conducted in semi-solid (0.3%) agar plates containing 10% NB in M9 medium plates. Mutants from the Keio collection, Δ*ugd* and Δ*wzc*, were screened for their swarming efficiencies and compared with wild type, *manA* and *wcaG* mutants, these latter two mutants being defective for capsular polysaccharide synthesis. The swarming rates detected for the *ugd* and *wzc* mutants increased around 3-fold (Table 2), while no change in swarming rate was observed for mutants incapable of incorporating fucose or mannose into LPS during its biosynthesis. These mutants were Δ*wcaG*, Δ*manA* and Δ*uxaC*. Thus, the interaction between CheY and Ugd-P has been confirmed, and this interaction is physiologically important for reciprocal regulation of motility and extracellular polysaccharide synthesis in response to stressful environments.

### Two-hybrid bacterial assay

CheY-Ugd and RssB-Ugd interactions were confirmed using two-hybrid assays after one day of bacterial growth at 30°C, followed by subsequent entry into the stationary growth phase at room temperature. At 37° C, only weak interactions were observed between CheY and Ugd.

## Discussion

Ugd is involved in lipopolysaccharide modification and CA production in *E. coli*, known previously to be subject to allosteric regulation (see Introduction). In this paper, we report different types of regulation of Ugd activity, resulting from protein-protein interactions with other enzymes and CheY. The Ugd interaction with the regulator of flagella rotation, CheY, apparently results in reciprocal regulation of swarming versus CA biosynthesis.

WecC-dependent activation of Ugd activity may be functionally related to Ugd-dependent Lipid A modification (13,22). Possibly in response to the PmrAB-dependent overexpression of a cluster of Lipid A and exopolysacharide biosynthetic genes, Ugd is activated by WecC to incorporate glucuronate moieties into EPS (CA). Gmd-dependent activation of Ugd is probably essential when CA biosynthesis is induced. ATP-dependent Wzc-catalyzed phosphorylation of Ugd to give Ugd-P is important under several stress conditions (2,10). But under conditions where the ATP concentration is low, Gmd may activate Ugd by a direct protein-protein interaction mechanism.

Based on the results reported here, we believe that Ugd may play a central regulatory role, influencing several physiological processes in addition to the reciprocally regulated capsular polysaccharide (CA) production and motility. Phosphorylation of Ugd appears to have a reciprocal effect on the CheY protein, influencing cell swarming. We found that the phosphorylated form of Ugd (Ugd-P) probably influences motility, since the Δ*ugd* mutant swarming rate increased more than 2-fold relative to the wild type strain, and overexpression of *ugd* decreased motility equally dramatically. Although the loss of other genes involved in CA biosynthesis were without effect, a *wzc* deletion mutation increased the swarming rate to the same extent as the *ugd* mutant, suggesting that only Ugd-P inhibits swarming. Capsular polysaccharide synthesis is disrupted in *wcaG* and *manA* mutants, but no effect of these mutations on motility was detected, as also shown in (23). These results suggest that Ugd is a global regulator of swarming, possibly due to a direct interaction with CheY.

Capsular PSs are produced in small quantities under normal growth conditions at temperatures higher than 30°C, but they are essential under certain stressful growth conditions (24), such as those that occur at low temperatures, low pH, antibiotic stress or exposure to the gastrointestinal tracts of animals (25). CA is related to persistence in the environment for enteroaggregative *E. coli* virulent strains (23,26). Swarming is often beneficial under condition of nutrient limitation, but not in stationary biofilms, leading to an explanation for the reciprocal regulation of motility and capsular polysaccharide synthesis. Thus, competitive environments may lead to the evolution of protein conformations promoting interactions in response to different environmental signals such as low Mg^2+^, low Fe^2+^, osmotic stress, and the presence of polymyxin. In this regard, the responses of *E. coli* to environmental changes seem to be remarkably flexible, involving the complex Ugd-dependent regulatory circuits described for the first time in this communication.

The effect of Ugd loss or overexpression on swarming could be a consequence of a disturbance in the RssB-RpoS interaction. Like Ugd, RssB is controlled by tyrosyl phosphorylation. There are two kinases, one associated with the colanic acid exporter noted above, and another activated by stress conditions. When RssB is phosphorylated, it binds RpoS and feeds it to proteasomes for degradation (27). This happens during log phase growth, but not during the stationary growth phase. This process is influenced by IraD, IraM and IraP, each which responds to different stress signals occurring during the stationary growth phase (27). Ugd interacts with RssB, and also inhibits swarming, while deletion of *ugd* causes an increase in swarming, but how? Maybe swarming is additionally regulated through RpoS and RssB. Further studies will be required to elucidate the precise mechanisms involved.

## Acknowledgements

We thank Dr. Granasse for the gift of the *wzc* construct, allowing overproduction of the soluble Wzc kinase domain and the achievement of a high degree of its purity. This work was supported by NIH grant GM109895. We thank Cali Myers and Harry Zhou for help in the preparation and submission of this manuscript.

